# TERSE: Efficient compression of the diffraction data

**DOI:** 10.1101/2023.05.10.540139

**Authors:** Senik Matinyan, Jan Pieter Abrahams

## Abstract

High-throughput data collection in crystallography poses significant challenges in handling massive amounts of data. Here, we present TERSE, a novel lossless compression algorithm specifically designed for diffraction data. We compare TERSE with the established lossless compression algorithms implemented in gzip, CBF, and HDF5, in terms of compression efficiency and speed, using continuous rotation electron diffraction data of an inorganic compound. Our results show that TERSE outperforms these algorithms by achieving a higher data compression at a speed that is at least an order of magnitude faster. TERSE files are byte-order independent and the algorithm can be readily implemented in hardware. By providing a tailored solution for diffraction data, TERSE facilitates more efficient data analysis and interpretation while mitigating storage and transmission concerns. TERSE C++20 compression/decompression code and an ImageJ/Fiji java plugin for reading TERSE files are open-sourced on GitHub under the permissive MIT license.

**Synopsis:** We present a fast and lossless algorithm for compressing diffraction data, achieving up to 85% reduction in file size while processing up to 2000 512×512 frames per second. This breakthrough in compression technology is a significant step towards more efficient analysis and storage of large diffraction datasets.

## 1. Introduction

Universal access to exponentially growing data has made efficient data storage and processing crucial for transformative science (Hill *et al*., 2016; Tolle *et al*., 2011). In crystallography, the emergence of hybrid pixel detector technology has led to a significant increase in the amount of data generated per data collection session, producing exponentially growing volumes of diffraction data (Paton *et al*., 2021; Tate *et al*., 2016). Because these detectors are so fast and have no readout-noise, fine phi-slicing and high frame rates allow more accurate data, with many pixels having values close to zero. However, the resulting high acquisition rates of diffraction data are outpacing the data transfer capabilities to local storage, presenting a significant challenge (Kieffer *et al*., 2018; Stroppa *et al*., 2023). Additionally, the size of current datasets poses challenges for transferring, sharing, and collaborating effectively, leading to increased operational costs, reduced experimental throughput, and potentially lost scientific information due to inefficiencies in data handling.

To address these challenges, there is an urgent need for robust, more efficient diffraction data compression that is lossless and fast enough to keep up with the high frame rates of modern detectors. In this context, we present the TERSE algorithm, a novel compression method specifically designed for diffraction data. Initial tests have shown that TERSE can compress integral data up to 15% of its initial size and handle up to 2000 electron diffraction frames per second (512×512 16-bit pixels), using a modern laptop. By providing a tailored solution for diffraction data, TERSE mitigates storage and transmission concerns. This algorithm has the potential to significantly improve the handling and long-term preservation of high-throughput diffraction data, facilitating scientific discoveries and accelerating the pace of transformative science.

## 2. TERSE Compression Algorithm

Diffraction data frames typically consist of a large number of grayscale pixels with integral values and a high dynamic range. At lower resolutions, the pixels tend to have higher values in Bragg peaks, while between the Bragg peaks and at higher resolutions, they have lower values. Thus, diffraction data frames are spatially correlated. By leveraging these inherent properties, TERSE performs lossless, efficient, and fast compression of integral diffraction data frames and other integral grayscale data. The algorithm was specifically designed for speed, but we found it to be also superior in reducing file sizes.

### 2.1. Compression Scheme

The TERSE algorithm uses a run-length encoding approach, which compresses data by identifying repeated patterns (Robinson & Cherry, 1967). Unlike most other run-length encoding algorithms, it requires just a single pass through the data, analogous to an early algorithm for compressing diffraction data (Abrahams, 1993). Implemented in C++20, it can easily be linked into other programs, that may be written in computer languages than C++20. The code creates a ‘Terse’ object using either an standard data container, a memory location, or a stream of raw data with a pixel depth of up to 64 bits. The algorithm compresses data quickly and efficiently by identifying patterns in a single pass using primitive processor operators. The resulting Terse object can be written as a byte-stream, which is independent of the endianness of the machine, ensuring that both big- and little-endian machines produce identical files. Because it can also be appended to existing files, it can be embedded in other data formats that may have specific header information, by replacing the raw data section. A Terse file has a small XML header that contains essential metadata required for unpacking, and that can easily be extended for specific use cases. The binary Terse data directly follows the XML header. For positive data, compressing as an unsigned integer yields a tighter compression. For data with negative integral numbers, TERSE uses 2-complement format for encoding, where the negative number is represented by the two’s complement of its absolute value.

Files that contain TERSE data can be read directly into a Terse object, which can be decoded by its Terse::prolix() member function. The Terse::prolix() member function allows the user to specify the location where the unpacked data will be stored, by providing an iterator or memory location as the argument. A Terse object can be unpacked into any type of arithmetic data, including floats and doubles. However, when the data include negative values, these cannot be unpacked into unsigned integers, and also data cannot be unpacked into integers with fewer bits than the original data.

#### 2.1.1. Block compression

TERSE compresses the data in fixed-size blocks (Figure 1). The pixel values of each data block (by default 12 integral values) are stripped of their most significant bits, provided they are all either zero (for unsigned values), or all identical (for signed values). In the latter case, the sign bit is maintained.

**Figure 1.**
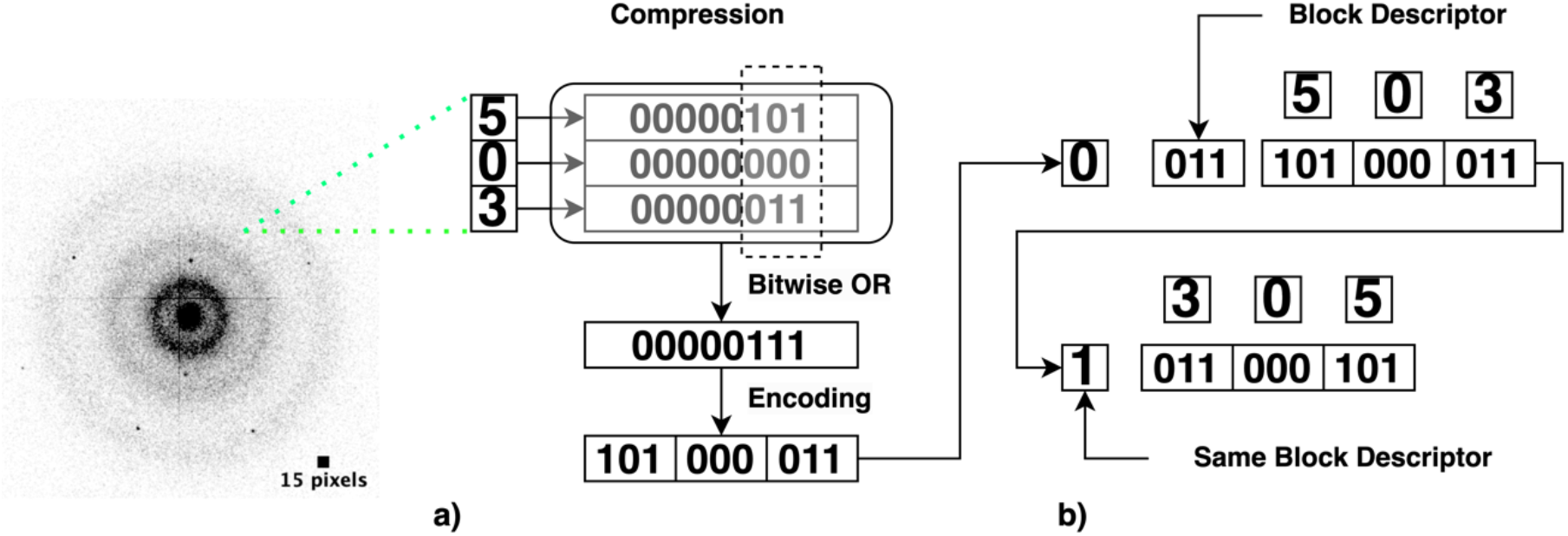
Compression scheme. a) For a block size of 3 with values 5, 0, 3, the encoded bits would be: 101(denoting 5) 000(denoting 0) 011(denoting 3). This block of data can therefore be encoded as 3 values of 3 bits each. The encoded 101000011would be pushed into the Terse object. b) Each block descriptor is preceded by a single bit. If the bit is set, the block descriptor is identical to the previous one. If the bit is not set, a new block descriptor follows. In this scenario, bits 2 to 4 define how many bits are used per value of the encoded block. If all three subsequent bits are set, the block descriptor is expanded to allow encoding pixel values that require up to 64 bits.

Each compressed data block is preceded by a variable-sized block descriptor indicating the number of bits used for encoding a single pixel value. In diffraction data, the required number of bits can vary from 0 to 64. However, lower values (requiring 0 to 6 bits, corresponding to a dynamic range of 0 to 128) are more common than higher values (requiring 10 to 64 bits, corresponding to a dynamic range of 2048 to 1.8 − 10^19).

To optimize compression, the block descriptor has a length of either 1, 4, 6, or 12 bits. The structure of the block descriptor is as follows:

Bit 1: If set, the previous block descriptor is uper second using a single CPU core on a modern laptop. Withsed; if not, the descriptor is expanded with 3 more bits.

Bits 2 to 4: These indicate how many bits are used per pixel value in the encoded block. If all three bits are set, then 7 or more bits per value are required, and the descriptor is expanded with an additional 2 bits.

Bits 5 and 6: The first 4 header bits must be 0111. The number encoded by bits 5 and 6 is added to decimal 7 to determine the number of bits used to encode each value in the block. Specifically, if bits 5 and 6 are 00, then 7 bits are used; if they are 01, then 8 bits are used; if they are 10, then 9 bits are used; and if they are 11, then at least 10 bits are used. If both bits 5 and 6 are set, the header is expanded by an additional 6 bits.

Bits 7 to 12: The first 6 header bits must be 011111. The number encoded by bits 7 to 12 is added to decimal 10 to determine the total number of bits used to encode each value in the block. Specifically, if bits 7 to 12 are 000000, then 10 bits are used; if they are 110110, then 64 bits are used (i.e., 10 + 54).

While other encoding schemes are possible, this particular one was found to be optimal for weak diffraction data and virtually indistinguishable from others for strong diffraction data. By using a variable-sized block descriptor and allowing for identical descriptors to be used for adjacent blocks, the encoding scheme can efficiently compress the data while retaining essential information.

## 3. Comparative Analysis of Terse Compression Algorithm with gzip, CBF and HDF5 with LZF and gzip Filters

### 3.1. Test Dataset

Continuous rotation electron diffraction data of an inorganic crystal were collected at PSI (Villigen, Switzerland). The diffraction experiment was carried out using a JEOL F200 TEM with a Schottky field emission gun (FEG) and a CEOS CEFID energy filter, and was operated at 200 keV. The detector used was an ASI Cheetah M3 retractable hybrid pixel detector, which collected zero-loss data as 16-bit 512 × 512 pixel .tiff stacks. The data were acquired in continuous, low gain mode at a rate of 10 frames per second while the sample was continuously rotated at 1.4 °/s. The unstacking of the data was performed using the EMAN2 package (Tang *et al*., 2007), resulting in 450 frames of 16-bit 512 × 512 pixel data. The data take up 237.8 MB of disk space.

### 3.2. Results

We evaluated the performance of several compression algorithms, including TERSE, gzip (compression level = 6), Crystallographic Binary File (CBF), and HDF5 with LZF and gzip lossless compression filters. Our evaluation was based on several metrics, including compression rate, compression and decompression speeds, and CPU utilization. Our results, obtained using a MacBook Pro with an M1 Max processor using a single core in the case of TERSE, and averaged over five cycles, are summarized below:

- TERSE:The TERSE algorithm reduced the data size to 38.1 MB after compression, which corresponds to 84.0% compression efficiency. The compression speed was 0.22s user time for 450 frames, with a moderate 46% CPU utilization. The decompression speed was also fast at 0.17s user time with a 50% CPU utilization.
- gzip: gzip compressed the data to 48.4 MB, which corresponds to 79.6% compression efficiency. However, the compression speed was relatively slow at 15.6s user time, with a high CPU utilization of 87%. The decompression speed was 1.29s user time with a CPU utilization of 68%. When using gzip with the maximum compression level of 9, the resulting compressed dataset size was reduced to 46.2 MB. However, this increased compression level came at the cost of a significantly longer processing time.
- CBF: The CBF algorithm reduced the data size to 119.8 MB after compression, which corresponds to 49.6% compression efficiency. The compression speed was relatively slow at 2.68s user time, with a high CPU utilization of 92%. The decompression speed was 4.72s user time with a lower CPU utilization of 52%.
- HDF5 with LZF compression filter: The HDF5 format with LZF compression filter reduced the data size to 95.2 MB, which corresponds to 59.9% compression efficiency. The compression speed was 2.64s user time with a moderate CPU utilization of 40%, while the decompression speed was 1.32s user time, with a CPU utilization of 67%.
- HDF5 with gzip compression filter: The HDF5 format with gzip compression filter compressed the data to 50.9 MB, which corresponds to 78.6% compression efficiency. The compression speed was 4.1s user time with a moderate CPU utilization of 54%, while the decompression speed was 1.46s user time, with a CPU utilization of 66%.

#### 3.2.1 In-memory Compression and Decompression Performance

For the evaluation of in-memory compression and decompression performance, 450 frames were loaded into memory to eliminate I/O overhead from the calculations. The compression and decompression times were then measured and averaged over 20 cycles for each algorithm (Table 1). TERSE had the fastest compression and decompression speeds, followed by HDF5/LZF. Gzip and CBF had moderate performance in both compression and decompression times.

**Table 1.** Compression efficiency is calculated as the percentage reduction in file size compared to the initial size. CPU utilization refers to the percentage of CPU resources utilized during compression or decompression. Compression and decompression speeds refer to ‘user time’, and exclude ‘system time’ for program initialization, memory management and I/O. (De)compression speed without I/O overhead is the ‘wall clock time’ required for in-memory (de)compression. All tests were performed on the same hardware and software setup.

| Algorithm | Compression Efficiency (%) | Compression Speed (s) | Compression CPU Utilization (%) | Decompression Speed (s) | Decompression CPU Utilization (%) | Compression speed without I/O overhead (s) | Decompression speed without I/O overhead (s) |
| --- | --- | --- | --- | --- | --- | --- | --- |
| TERSE | 84.0 | 0.22 | 46 | 0.17 | 50 | 0.19 | 0.14 |
| gzip | 79.6 | 15.6 | 87 | 1.29 | 68 | 13.47 | 0.53 |
| CBF | 49.6 | 2.68 | 92 | 4.72 | 52 | 0.6 | 2.27 |
| HDF5/LZF | 59.9 | 2.64 | 40 | 1.32 | 67 | 0.68 | 0.407 |
| HDF5/gzip | 78.6 | 4.1 | 54 | 1.46 | 66 | 3.34 | 0.73 |

Based on the benchmark results, it is clear that the TERSE compression algorithm outperforms other compression algorithms such as gzip, CBF, and HDF5 with LZF and gzip compression filters. TERSE achieves a significantly higher data reduction rate while maintaining faster compression and decompression speeds. These results demonstrate the efficiency and effectiveness of the TERSE algorithm for compressing diffraction data, making it a valuable tool for handling large datasets in crystallography and other fields.

## 4. Discussion

The increasing availability of high-throughput data collection techniques in structural biology has created challenges in handling the massive amounts of data (Mokso *et al*., 2017). Different compression methods have been developed to address these challenges, such as compressing signals with singularities and transient phenomena, exploiting ptychographic oversampling, or reducing data based on simple azimuthal regrouping (Ferrer *et al*., 1998; Loetgering *et al*., 2017; Kieffer *et al*., 2018)

In this paper, we introduced the TERSE algorithm, a compression method specifically designed for diffraction data. The TERSE algorithm outperforms conventional compression techniques, such as gzip and CBF, in terms of both compression efficiency and speed. Our initial tests show that TERSE can almost keep up with up a data stream of up to about 2000 frames (512×512 pixels) per second using a single CPU core on a modern laptop. With extra hardware and parallel processing, it can potentially keep up with the fastest detectors available.

The generated TERSE (.trs) files are byte-order independent, which ensures compatibility across different hardware architectures. Furthermore, the compiled TERSE algorithm has a very small footprint and can be readily implemented in hardware, such as FPGA or ASIC, allowing for potential integration with existing data acquisition systems. This integration can greatly enhance the real-time processing capabilities and overall efficiency of data collection and storage in structural biology experiments.

By providing a tailored solution to handle the specific requirements of diffraction data, the TERSE algorithm not only mitigates storage and transmission concerns but also facilitates more efficient data analysis and interpretation. It aims to improve the efficiency of data storage and transmission, while retaining the essential information within the diffraction data.

In conclusion, further development and widespread adoption of the TERSE algorithm, and its integration with user-friendly data processing and analysis tools has the potential to streamline the workflow for researchers working with high-throughput diffraction data. This can ultimately accelerate research.

## Acknowledgements

This project has received funding from the European Union’s Horizon 2020 research and innovation programme under the Marie Skłodowska Curie grant agreement #956099, and from the Swiss National Science Foundation project grant #205320_201012. We would like to acknowledge Dmitry Byelov, Rick Watertor, and Ciaran Welsh from Amsterdam Scientific Instruments for their substantial contributions to the testing and debugging of the algorithm.

## Appendix A. Data and Code Availability

The TERSE compression algorithm, the associated TIFF library, and the ImageJ/Fiji plugin (Appendix B) are available in our GitHub repository (https://github.com/Senikm/terse.git) under the MIT License. This permissive open-source license allows for the free use, modification, and distribution of the software, with minimal restrictions, enabling the scientific community and other interested parties to build upon and integrate these resources into their own projects or workflows.

## Appendix B. Supplementary Materials

### B1.1. Terse C++20 code

With the open-source C++ header library, integral data stored in any memory location can be compressed in memory and subsequently be written to disk. Also, compressed data can be decompressed and read into any type of C++ container or memory location, storing any type of numerical values. All code is implemented in templated headers, with one header file containing the Terse-object code and two header files for XML parsing and bit manipulations.

### B1.2. TIFF Library

As an example of usage, the distributed package also contains C++20 code for compressing and decompressing TIFF files produced by Medipix Quad detectors, that have the following characteristics:

- one image per TIFF file;
- one grayscale intensity value per pixel;
- the intensity data always begin on position *8* in the TIFF file, to allow them being read as raw data by programs without TIFF libraries;

#### Parts that can be overridden

- by default, the intensities are stored in the file 16-bit unsigned integers
- by default, the image size is 512^*^512 pixels.

Compressed files have a .trs extension.

### B1.3. ImageJ Plugin for TERSE (.trs) Format Data

The Terse Reader plugin is designed for reading, unpacking, and visualizing image data from .trs files in ImageJ/Fiji. The plugin reads the image data from a selected .trs file and displays it as a 512^*^512-pixel grayscale image. When executed, the plugin first prompts the user to select the data file. Once the file is selected, the plugin reads its XML header to obtain important parameters required for unpacking the .trs file. Once unpacked, the plugin creates an object with dimensions 512^*^512, populates the object with the unpacked data, and displays the resulting image.

## References

Abrahams, J. P. (1993). Joint CCP4 and ESF-EACBM Newsletter on Protein Crystallography 28, 3–4.

Ferrer, J. L., Roth, M. & Antoniadis, A. (1998). Acta Crystallogr D Biol Crystallogr 54, 184–199.

Hill, J., Mulholland, G., Persson, K., Seshadri, R., Wolverton, C. & Meredig, B. (2016). MRS Bulletin 41, 399–409.

Kieffer, J., Petitdemange, S. & Vincent, T. (2018). J Synchrotron Rad 25, 612–617.

Loetgering, L., Rose, M., Treffer, D., Vartanyants, I. A., Rosenhahn, A. & Wilhein, T. (2017). Advanced Optical Technologies 6, 475–483.

Mokso, R., Schlepütz, C. M., Theidel, G., Billich, H., Schmid, E., Celcer, T., Mikuljan, G., Sala, L., Marone, F., Schlumpf, N. & Stampanoni, M. (2017). J Synchrotron Rad 24, 1250–1259.

Paton, K. A., Veale, M. C., Mu, X., Allen, C. S., Maneuski, D., Kübel, C., O’Shea, V., Kirkland, A. I. & McGrouther, D. (2021). Ultramicroscopy 227, 113298.

Robinson, A. H. & Cherry, C. (1967). Proceedings of the IEEE 55, 356–364.

Stroppa, D. G., Meffert, M., Hoermann, C., Zambon, P., Bachevskaya, D., Remigy, H., Schulze-Briese, C. & Piazza, L. (2023). Microscopy Today 31, 10–14.

Tang, G., Peng, L., Baldwin, P. R., Mann, D. S., Jiang, W., Rees, I. & Ludtke, S. J. (2007). Journal of Structural Biology 157, 38–46.

Tate, M. W., Purohit, P., Chamberlain, D., Nguyen, K. X., Hovden, R., Chang, C. S., Deb, P., Turgut, E., Heron, J. T., Schlom, D. G., Ralph, D. C., Fuchs, G. D., Shanks, K. S., Philipp, H. T., Muller, D. A. & Gruner, S. M. (2016). Microscopy and Microanalysis 22, 237–249.

Tolle, K. M., Tansley, D. S. W. & Hey, A. J. G. (2011). Proceedings of the IEEE 99, 1334–1337.

